# Asymmetric compression of representational space for object animacy categorization under degraded viewing conditions

**DOI:** 10.1101/166660

**Authors:** Tijl Grootswagers, J. Brendan Ritchie, Susan G. Wardle, Andrew Heathcote, Thomas A. Carlson

**Affiliations:** Macquarie University, Australia; ARC Centre of Excellence in Cognition and its Disorders, Australia; University of Sydney, Australia; KU Leuven, Belgium; University of Tasmania, Australia; University of Newcastle, Australia

## Abstract

Animacy is a robust organizing principle amongst object category representations in the human brain. Using multivariate pattern analysis methods (MVPA), it has been shown that distance to the decision boundary of a classifier trained to discriminate neural activation patterns for animate and inanimate objects correlates with observer reaction times for the same animacy categorization task (Carlson, Ritchie, Kriegeskorte, Durvasula, & Ma, 2014; Ritchie, Tovar, & Carlson, 2015). Using MEG decoding, we tested if the same relationship holds when a stimulus manipulation (degradation) increases task difficulty, which we predicted would systematically decrease the distance of activation patterns from the decision boundary, and increase reaction times. In addition, we tested whether distance to the classifier boundary correlates with drift rates in the Linear Ballistic Accumulator (Brown & Heathcote, 2008). We found that distance to the classifier boundary correlated with reaction time, accuracy, and drift rates in an animacy categorization task. Split by animacy, the correlations between brain and behavior were sustained for longer over the time course for animate than for inanimate stimuli. Interestingly, when examining the distance to the classifier boundary during the peak correlation between brain and behavior, we found that only degraded versions of animate, but not inanimate, objects had systematically shifted towards the classifier decision boundary as predicted. Our results support an asymmetry in the representation of animate and inanimate object categories in the human brain.

## 1. Introduction

Object recognition is a fast, reliable, and effortless process for humans. Early visual areas in the brain respond to simple visual features (e.g., edges, luminance contrast, or orientation), and further along the ventral stream sensitivity to objects and object categories (e.g., faces, animals, or tools) emerges (DiCarlo & Cox, 2007; Grill-Spector & Weiner, 2014). Using multivariate pattern analysis (MVPA), several studies have analyzed pattern similarities between the neural representation of objects in inferior temporal cortex (ITC) to study its representational structure, with both fMRI (Edelman, Grill-Spector, Kushnir, & Malach, 1998; Haxby, Connolly, & Guntupalli, 2014; Kriegeskorte et al., 2008; Kriegeskorte & Kievit, 2013), and MEG (Carlson, Tovar, Alink, & Kriegeskorte, 2013; Cichy, Pantazis, & Oliva, 2014, 2016; Contini, Wardle, & Carlson, in press). These studies have provided evidence of a categorical organization in ITC, following the observation that objects belonging to the same category tend to evoke similar patterns of neural activation (Connolly et al., 2012; Haxby et al., 2001; Kriegeskorte et al., 2008; Sha et al., 2015). Time series decoding studies have shown that such abstract object representations emerge later than simple visual features in the time-course of object processing (Carlson et al., 2013; Cichy et al., 2014, 2016; Kaneshiro, Guimaraes, Kim, Norcia, & Suppes, 2015), consistent with a hierarchical organization of the visual stream. One robust categorical structure is the animate/inanimate distinction in human and primate ITC (Caramazza & Shelton, 1998; Kiani, Esteky, Mirpour, & Tanaka, 2007; Kriegeskorte et al., 2008; Mahon & Caramazza, 2011; Spelke, Phillips, & Woodward, 1995). Animate stimuli (e.g., humans or animals) evoke brain activation patterns that are more similar to other animate exemplars than to inanimate stimuli (e.g., plants, tools, vehicles) (Carlson et al., 2013; Cichy et al., 2014; Downing, Jiang, Shuman, & Kanwisher, 2001; Kiani et al., 2007; Kriegeskorte et al., 2008).

A current topic of debate is whether categorical information in the brain, as revealed using MVPA decoding, is read-out in behavior (cf. de-Wit, Alexander, Ekroll, & Wagemans, 2016; Williams, Dang, & Kanwisher, 2007). For the presence of decodable information in neuroimaging activation patterns related to a stimulus or task does not necessarily entail that this information underlies related behavior. One recent approach to linking activation spaces to behavior (Ritchie & Carlson, 2016) is inspired by distance-to-bound models of reaction time (Ashby & Maddox, 1994; Pike, 1973). According to distance-to-bound models, evidence close to a decision boundary is more ambiguous, reflecting greater difficulty in categorization, while evidence far from the decision boundary is less ambiguous with regards to category membership. Assuming that response time is a function of stimulus discriminability, and that a classifier decision boundary (which is used in MVPA decoding) reflects an observer’s decision boundary, then reaction times should negatively correlate with distance from the boundary; for example, stimuli that are faster to categorize should be neurally represented as further from the classifier decision boundary (Ritchie & Carlson, 2016).

In a previous application of the RT-distance approach to MEG (Ritchie et al., 2015), a linear discriminant classifier (LDA) was trained to discriminate the MEG channel activation for animate from inanimate stimuli. Next, the distance of each stimulus pattern to the classifier boundary in high-dimensional space were rank-order correlated to human reaction times for categorizing the same stimuli as animate/inanimate. As predicted, the distance to boundary negatively correlated with reaction time. Moreover, the correlation over time tracked the MEG decoding time-series (Ritchie et al., 2015). Interestingly, when analyzing the animate and inanimate stimuli separately, an asymmetry was observed: the correlation between distance to boundary and reaction time was driven by the animate stimuli (Carlson et al., 2014; Ritchie et al., 2015).

A useful test of the RT-distance hypothesis would be to manipulate task difficulty experimentally, and observe the effects of this behavioral manipulation on representational space. Note that here we use the term ‘representational space’ under the assumption that neural representations are to some extent reflected in neuroimaging activation patterns (see e.g., Carlson, Schrater, & He, 2003; Haxby et al., 2001; Kriegeskorte & Bandettini, 2007). The RT-distance hypothesis has so far been tested on the differences *between* objects. For example, in Carlson et al, (2014), an ostrich was closer to the animacy decision boundary than a human face, and participants were slower to categorize the ostrich as ‘animate’. The effect of increasing categorization difficulty of single object images on the distance to the classifier boundary in activation space has not yet been tested. According to the RT-distance hypothesis, a change in behavior resulting from manipulating categorization task difficulty should be matched by a corresponding shift of the stimulus set in representational space. Numerous studies have shown that degrading object stimuli reduces categorization performance, such as by scrambling the image phase (e.g., Philiastides, Ratcliff, & Sajda, 2006; Philiastides & Sajda, 2006; Wichmann, Braun, & Gegenfurtner, 2006), scrambling image amplitude (e.g., Gaspar & Rousselet, 2009), reducing luminance contrast (e.g., Macé, Delorme, Richard, & Fabre-Thorpe, 2009; Macé, Thorpe, & Fabre-Thorpe, 2005), or blurring the image (e.g., Bruner & Potter, 1964; Párraga, Troscianko, & Tolhurst, 2000, 2005; Wyatt & Campbell, 1951). For degraded stimuli with longer categorization RTs, the RT-distance hypothesis predicts that these stimuli will be located closer to the animacy decision boundary, producing a ‘compression’ of representational space compared to that for the original versions of the stimuli (Fig 1A). This predicts a correlation between the shorter distance to boundary and slower accumulation rates (Fig 1B). Note that even in the case of inanimate stimuli where there was no correlation (Ritchie et al., 2015), one would still expect a correlation when including both clear and degraded versions because of the compression of the general representational space (Fig 1B).

**Figure 1.**
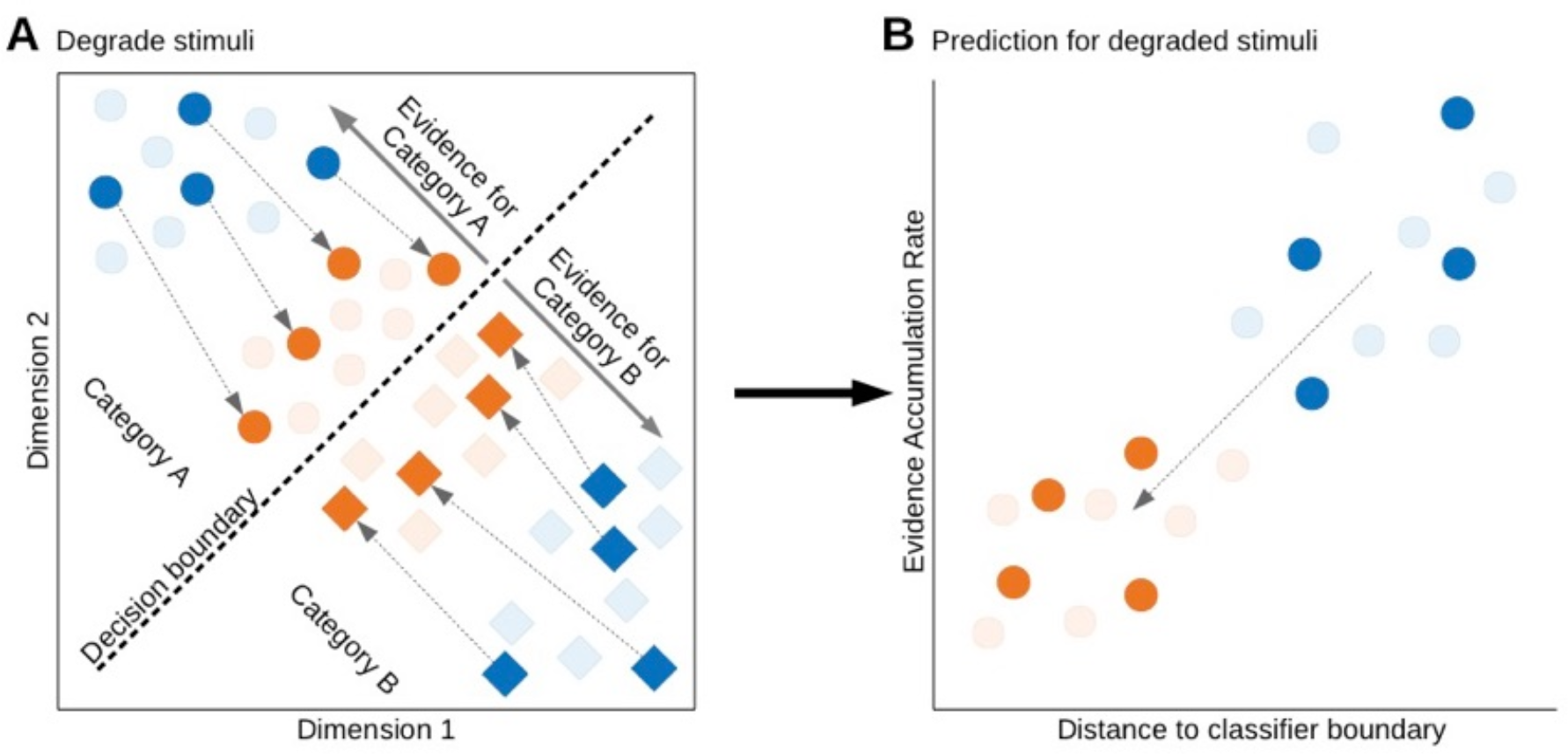
The predicted effect of degrading stimuli on their location in representational space. **A.** Stimuli of two categories are illustrated as circles (animate objects) and squares (inanimate objects) in representational space (only two dimensions are plotted here for visualization). The blue shapes represent the stimuli in a clear state, and the orange are their degraded counterparts. If distance to a classifier boundary is taken as representing evidence for a decision, this predicts that the degraded (orange) versions of the stimuli will be located closer to the classifier decision boundary (dashed line) that separates the stimulus categories than the clear (blue) versions. **B.** Stimuli from one category (animate objects) and their distance to the classifier boundary versus their rate of evidence accumulation for an animacy categorization decision. The RT-distance hypothesis predicts that the degraded stimuli, which have moved closer to the decision boundary, also have slower evidence accumulation rates (and therefore longer RTs). Note that the same prediction holds for the other category (inanimate objects).

The dominant models of reaction time appeal to some form of evidence accumulation process (Ashby, 2000; Ratcliff, 1985); That is, evidence for a decision (e.g., animate or inanimate) accumulates over time, and the response is made when the amount of evidence reaches a certain threshold (Brown & Heathcote, 2008; Gold & Shadlen, 2007; Ratcliff & Rouder, 1998). Distance to the classifier boundary can be linked to evidence accumulation. Carlson et al., (2014) simulated evidence accumulation with a sequential analysis model, using the distance to the boundary for each object exemplar as a proxy for evidence strength. They found that accumulation rate correlated with categorization RTs, providing support for the theoretical link between distance to boundary and evidence accumulation (Carlson et al., 2014; Ritchie et al., 2015). One way to build upon this would be to more closely relate distance to evidence accumulation; beyond correlating distance with median RTs, an existing model of evidence accumulation can be fit to subjects’ behavioral data, yielding independent model parameter estimates that can be correlated with distance to the boundary. As an accumulator model provides a more complete characterization of categorization behavior than average reaction times, it may provide a better measure to correlate to representational distances.

The aim of the present study was two-fold. First, using MEG decoding we sought to test the prediction that degrading object exemplar would compress the representational space (which we assume is reflected in the high dimensional MEG activation space). This would in turn correlate with slower animacy RTs compared to un-degraded, or clear versions of the same stimuli (Figure 1). Secondly, we aimed to test the RT-distance hypothesis in the context of an existing model of RT distributions and choice accuracy, the Linear Ballistic Accumulator (LBA; Brown & Heathcote, 2008), in order to evaluate whether distance to boundary can be related more directly to evidence-accumulation model parameters.

## 2. Methods

### 2.1 Participants

All participants gave informed consent in writing prior to the experiment. The study was conducted with the approval of the Macquarie University Human Research Ethics Committee. 100 participants were recruited on Amazons Mechanical Turk (MTurk) to determine the level of stimulus degradation needed for equal object recognition performance across all stimuli (see Section 2.1). Another group of 40 new MTurk participants were asked to rate the stimuli on a 7-point scale from typical animate to typical inanimate. For the second part of the study, 20 healthy volunteers (4 males; mean age 29.3 years) with normal or corrected-to-normal vision participated in the MEG experiment. Participants in both experiments were financially compensated for their time. All analysis procedures were performed in Matlab, using the statistics and machine learning toolbox. For reference purposes, the code is freely available at https://github.com/Tijl/Grootswagers_etal_degraded_objects.

### 2.1 Stimuli

We constructed a set of 48 visual object stimuli including 24 animals and 24 inanimate objects (natural and man-made) on a phase scrambled natural image background in both a clear and a degraded condition. First, high resolution images (> 512 × 512 pixels) of various objects were collected via an internet search. We only selected images that included prototypical viewpoints of objects. In order to test the RT-distance hypothesis more generally, no human or human face images were included in the stimulus set as they are generally outliers both behaviorally and in the brain’s response compared to other object exemplars. That is, face stimuli tend to have fast categorization RTs and produce pronounced responses in neuroimaging data. It is therefore possible that the inclusion of face stimuli could disproportionately explain any observed correlations between RT and neural distance. All exemplar images used in the study are shown in Figure 2A. As color can be a salient cue for image recognition (Biederman & Ju, 1988; Joseph & Proffitt, 1996; Ostergaard & Davidoff, 1985; Wurm, Legge, Isenberg, & Luebker, 1993) and would make the degrading process (described in the next section) less effective, gray-scaled versions of the object images were used in the experiment. A different random noise background was created for each exemplar by phase scrambling a natural image of a forest scene (see Figure 2B). The object images were overlaid on the noise background, producing the stimuli for the clear condition (Figure 2C).

**Figure 2.**
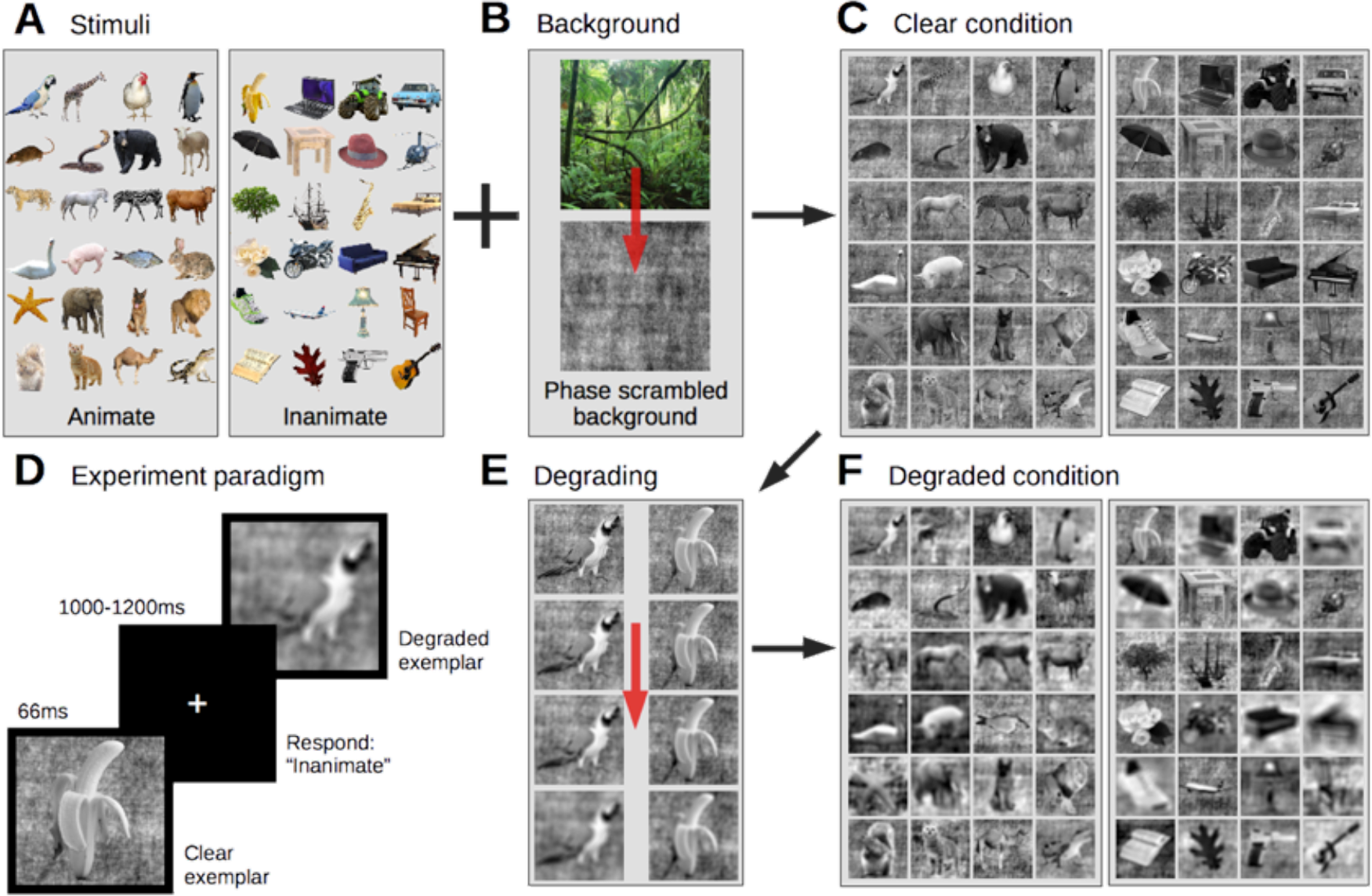
Stimuli and experiment design. **A.** Stimuli consisted of 24 animate and 24 inanimate objects. **B.** Stimuli were placed on a phase scrambled natural image background. **C.** All stimuli in the clear condition. **D.** In the MEG experiment, participants saw clear and degraded stimuli for 66ms in randomized order with a varying inter stimulus interval and were asked to report the stimulus category (animate or inanimate) with a button press. **E.** To create the degraded condition, stimuli were gradually blurred by simulating defocus to a level where they were equally recognizable. **F.** All stimuli in the degraded condition.

We created a degraded condition by blurring the same set of images (including their background). As different objects require different levels of blur to equate recognition performance, we first measured the amount of blur required to impair recognition for each image. Our aim was for subjects to be able to perform the task given unlimited exposure time (i.e., correctly recognize the object), but to reduce their categorization performance under brief presentation duration (i.e., by reducing speed and/or accuracy). We simulated defocused blur using an image filter (Figure 3) that convolved the amplitude spectrum of the image in the Fourier domain (Figure 3A-B) with a Fourier-transformed cylinder function (a sombrero function, Figure 3C-D), where increasing the radius of the cylinder function results in a greater magnitude (Figure 3E) of defocus blur (Sonka, Hlavác, & Boyle, 2008). Images were then gradually degraded by increasing the radius in steps of 2 pixels, from a radius of one pixel (no degrading) to 59 (very degraded). The sequence of the image coming into focus was presented to the MTurk participants. Each participant saw all 48 stimuli from both animate and inanimate categories once starting from the most degraded state, while its level of focus was gradually increased (in steps of 2 pixel radius). Participants were instructed to press the spacebar as soon as they recognized the object in the picture. The stimulus was then removed from the screen, and participants entered a name for the stimulus. To check for correct recognition, the responses on the naming task were assessed manually for validity, to allow for variations in spelling or for synonymous names. For each exemplar, the amount of focus needed for 25% of the MTurk participants to correctly recognize (i.e., name) the object (see Figure 2F) was used as the blur filter parameters for that exemplar in the degraded condition. On average, a radius of 17 pixels (sd=4.68) was used for the animate exemplars and 20.5 pixels (sd=10.12) for the inanimate exemplars.

**Figure 3.**
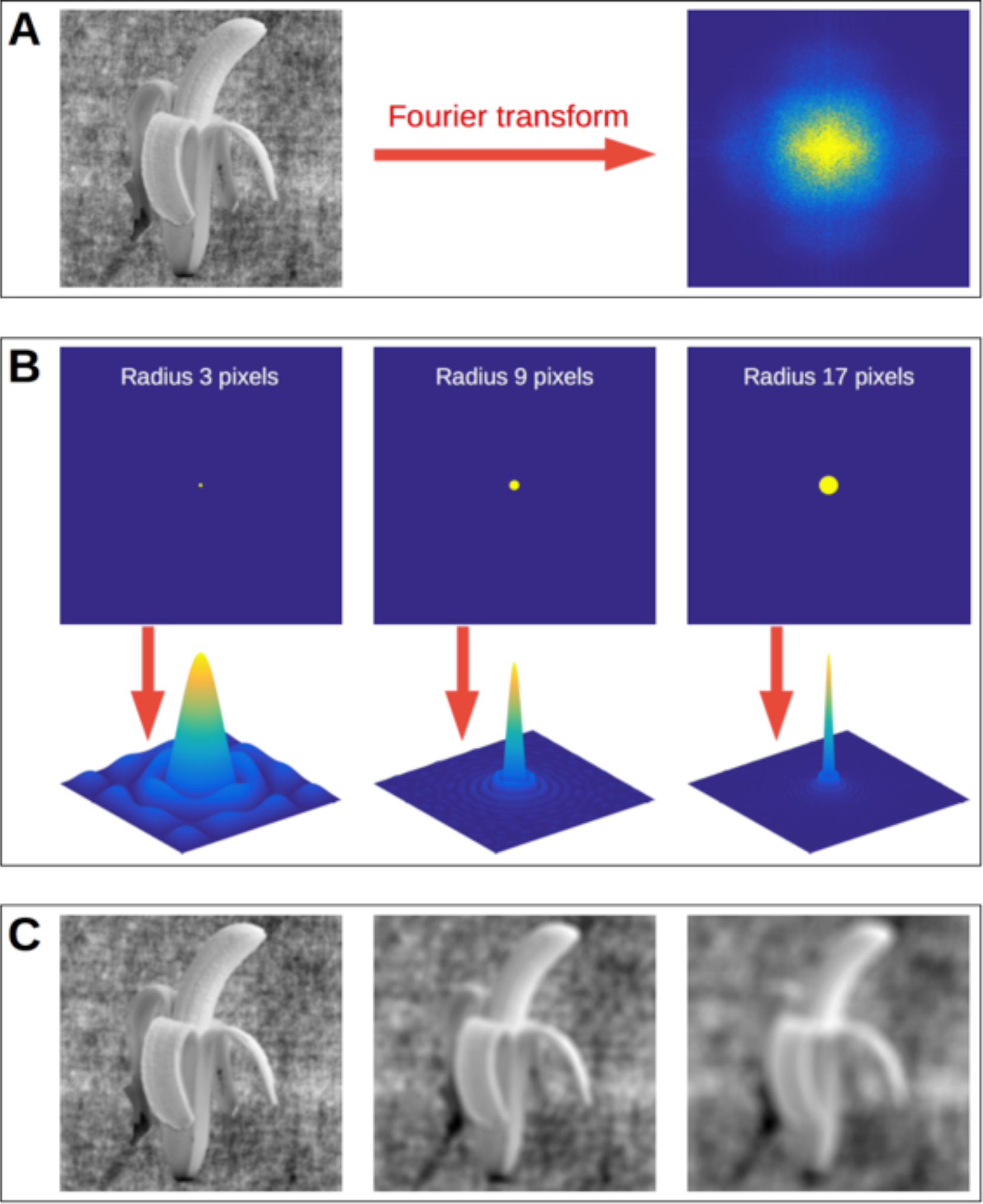
Image filter used to blur the experimental stimuli. **A.** The original image and its amplitude spectrum, which shows the typical energy pattern for natural images (high energy at low spatial frequencies (center), and low energy at high spatial frequencies Field, 1987). **B.** Cylinder functions with radii of 3, 9, and 17 pixels and their Fourier transformed versions (sombrero functions), which are used as the image filters. **C.** Convolving the amplitude spectrum of the original image with the sombrero functions in the Fourier domain results in images with different levels of defocus blur by removing a significant proportion of energy at high spatial frequencies.

### 2.2 MEG Experiment Design

Before the MEG experiment, we confirmed that each participant could recognize all exemplars given unlimited presentation time, even in the degraded state, as the intention of the design was to decrease categorization speed and accuracy without making the stimuli unrecognizable. Stimuli in the degraded state were shown and participants were asked to name the object in each picture. If the participant failed to correctly recognize the stimulus, it was shown in the clear state, to ensure that all objects were correctly recognized. Next, the participant was trained on the task (outside the MEG), as the brief presentation duration and fast-pace of the categorization task required practice to master. If performance (animacy categorization accuracy) in the first block was lower than 80%, the participant was shown all the stimuli again in both states, to identify degraded exemplars they were unable to recognize, and then practiced again on a second training block. All 20 participants performed above 80% correct after the familiarization step.

Following the practice task, participants completed the MEG experiment. On each trial within a block, stimuli were projected (at 9x9° visual angle) on a black background for 66ms, followed by a fixation cross for a random duration between 1000 and 1200ms. Participants were asked to categorize the stimulus as animate or inanimate as fast and accurate as possible, using a button press (see also Figure 2D). To avoid possible motor response confounds, the mapping of the response buttons alternated between blocks (Grootswagers, Wardle, & Carlson, 2017; Ritchie et al., 2015) and participants received feedback for the first 10 trials of a block (red or green cross), to ensure accurate mapping of the buttons. Within each block, four repetitions of each exemplar in both conditions were presented in randomized order. Block duration was ~7 minutes. After each block, participant received feedback on performance (mean accuracy and number of missed trials). Each participant completed eight blocks, resulting in 32 trials per exemplar, 768 trials per category and condition (animate/inanimate, clear/degraded), and 3072 trials total per participant. The total time in the scanner was about one hour, including breaks between blocks.

### 2.3 MEG acquisition and preprocessing

Participants were fitted with a cap with 5 marker coils to track head movement during the session. The MEG signal was continuously sampled at 1000Hz from 160 axial gradiometers using a whole-head MEG system (Model PQ1160R-N2, KIT, Kanazawa, Japan) while participants lay in a supine position inside a magnetically shielded room (Fujihara Co. Ltd., Tokyo, Japan). Recordings were filtered online between 0.03Hz and 200Hz. We examined the delay in stimulus onsets (Ramkumar, Jas, Pannasch, Hari, & Parkkonen, 2013) by comparing the photodiode responses with the stimulus onset triggers (sent by the experiment script), and found a highly consistent delay of 56.26ms, for which we conservatively corrected by shifting the onset triggers back by 56ms. Recordings were sliced into 700ms epochs (-100 – 600ms post-stimulus onset). The trials were downsampled to 200Hz (5ms resolution) and transformed using principal component analysis (PCA), where the components that accounted for 99% of the variance were retained to reduce the dimensionality of the data (mean 62.25 components, sd 12.33). Finally, to increase signal to noise, 4 trials of each exemplar were averaged into pseudo-trials (Grootswagers et al., 2017; Isik, Meyers, Leibo, & Poggio, 2014), leaving 8 pseudo-trials per exemplar in both conditions.

### 2.4 Sliding time window decoding

To investigate the decoding performance of animacy over time, sliding window Naïve Bayes classifiers were used on the pseudo-trials. To assess the difference in decoding performance between the clear and degraded conditions, three separate classifiers were used for decoding; one for each condition, and one for both conditions combined. At each 5ms interval *t*, the classifiers were trained and tested on a 25ms window (from *t*-25ms to *t*). The classifier performance was examined using leave-one-exemplar-out cross-validation (Carlson et al., 2013). In this method, the classifier is trained on the animacy of all-but-one exemplar, and tested on trials of the left out exemplar. This is repeated for each exemplar, and the mean decoding accuracy on the left out exemplars is used to assess generalization accuracy. Using this cross-validation method, the classifier has to generalize the *concept* of animacy, and cannot benefit from exploiting individual stimulus properties because the test exemplar is not in the training set (Carlson et al., 2013). We report the subject-averaged classifier accuracy over time, with significant above-chance accuracies assessed using a non-parametric Wilcoxon signed rank test. The False Discovery Rate (FDR) was used to control for false positives resulting from multiple comparisons.

### 2.5 Fitting LBA on individual subject behavior

One of the aims of this study was to use a more complex model of evidence accumulation, and to then correlate distance to boundary with the drift rate parameters of the model, as well as RT and accuracy. The Linear Ballistic Accumulator is a mathematically tractable yet simple and complete model of evidence accumulation (LBA; Brown & Heathcote, 2008). LBA simultaneously takes into account both the full distribution of RTs (i.e., the mean, variability and positive skew of the RT distribution, see Luce, 1986) and task accuracy. LBA differs from other well-known accumulation models (e.g., the diffusion decision model Ratcliff, 1978, or the leaky competitive accumulator, Usher & McClelland, 2001) by modeling the response alternatives (here: animate and inanimate) with separate independent accumulators that accrue evidence accumulation in a linear and ballistic manner. In particular, the LBA model assumes that between-trial variability in the amount of evidence required for a decision, and the rate at which it accumulates, dominate over any moment to moment variability in evidence during a trial, whereas the latter source of noise plays a greater role in alternative models. These properties make LBA analytically simple and relatively easy to apply, which is ideal for this study, where each stimulus is treated as a separate condition, meaning that there are many parameters to estimate, each based on relatively few data points. To simplify the fitting procedure, the stimuli for both categories were separated into 6 bins based on accuracy. Next, a set of progressively more complex parameterizations were fit stepwise using maximum likelihood estimation (Donkin, Brown, & Heathcote, 2011; Rae, Heathcote, Donkin, Averell, & Brown, 2014). The most complex model parameterization, which was used here for further analysis, included clear/degraded, animate/inanimate, and stimulus as factors. Hence, each stimulus in both clear and degraded conditions had a separate drift rate, which is important to note, as the goal was to correlate stimulus-specific distances with drift rates. Figure 4 shows that the LBA model provided a very accurate model of the effect of stimulus degradation and of differences between animate and inanimate images, both in terms of accuracy (Figure 4A) and the entire distribution of RTs (Figure 4B), and so provides a useful characterization of the participants’ behavior.

**Figure 4.**
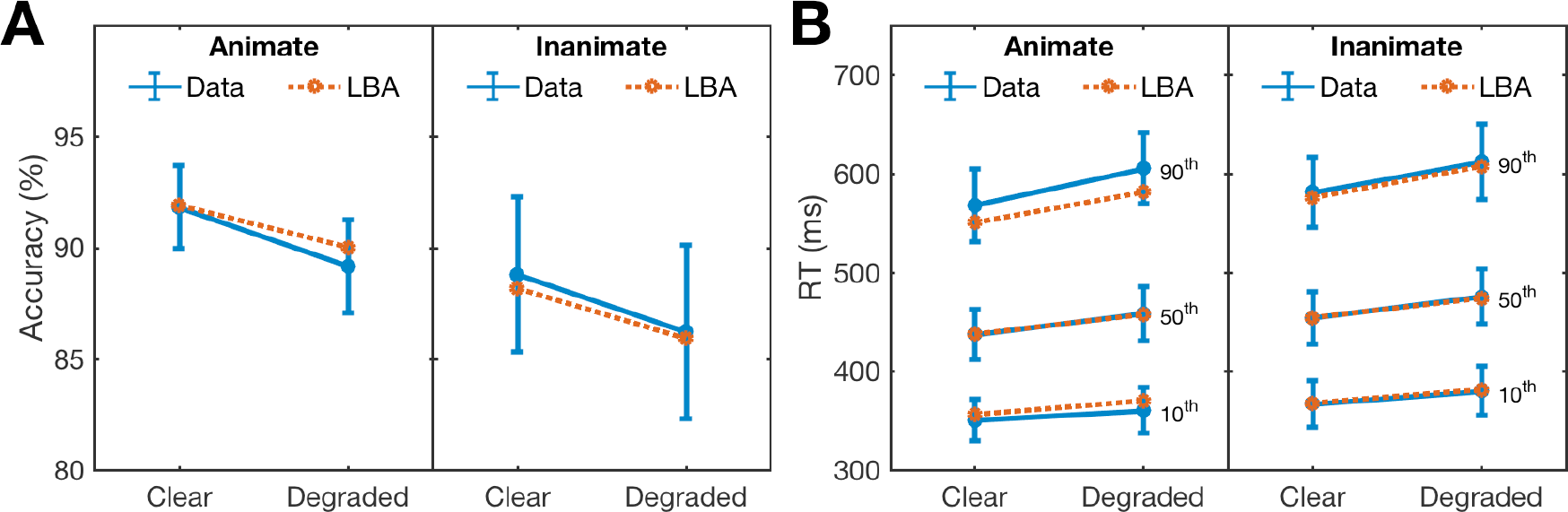
Fits of the LBA model to (A) accuracy and (B) the distribution of RT, with 95% confidence intervals. The RT distribution is illustrated by plotting the 90^th^ percentile (upper lines, representing the slowest RTs), the median (50^th^ percentile, middle lines) and the 10^th^ percentile (lower lines, representing the fastest RTs).

### 2.6 Predicting behavior from representational distance

Individual exemplars can be represented as points in representational space (i.e., a multidimensional feature space). To decode animacy, we applied a discriminant classifier (Gaussian Naïve Bayes) to optimize a decision boundary in multidimensional space, separating the neural patterns for animate and inanimate exemplars. According to the RT-distance hypothesis, the neural representations of the exemplars that are close (in multidimensional space) to this boundary are predicted to be more difficult to discriminate, as there is less evidence for the decision. This forms a prediction for behavior, where less evidence for a decision would result in slower RTs or lower accuracies (Carlson et al., 2014; Ritchie & Carlson, 2016; Ritchie et al., 2015), and, in the case of the LBA decision model, slower drift rates. For the current study, we predicted that degraded stimuli would be closer to the decision boundary in representational space, and therefore correlate with lower accuracy, slower reaction times, and slower drift rates.

We tested this prediction by repeating the following process at each time window: First, all the trials for each exemplar were averaged to create an average representation in multidimensional space for each exemplar (one for the exemplar in clear, and one for degraded state). In this space, a decision boundary for animacy was fitted (i.e., training a Naïve Bayes classifier), and the representational distance to this boundary for each exemplar was computed. This resulted in a distance-value for each exemplar in both clear and degraded states (24 animate and 24 inanimate exemplars in 2 states = 92 distance values). Next, the exemplar distances were rank-order correlated (using Spearman’s *ρ*) to the two behavioral measures (median reaction time, mean accuracy) and the mean drift rate for the exemplars.

Repeating this process over time and subjects resulted in three time-varying correlations (two for each behavioral measure, and one for drift rate) for each subject. We then report the subject-averaged time-varying correlations, and used a non-parametric Wilcoxon signed rank test at each time window to test for significant above-zero correlations at the group level. Note that this approach differs from Ritchie et al., (2015), where distances and reaction times were averaged over subjects first, and the correlations were performed at the group level. False discovery rate (FDR) adjustment was used to control for false positives resulting from multiple comparisons. This process allowed us to compare the distance to boundary correlations with the three variables. We also sought to assess the possibility of modulating effects of animacy, for example whether the animate-inanimate asymmetry reported in Carlson et al., (2014) and Ritchie et al. (2015) was replicated, and whether it was affected by degrading the stimuli. For this, we computed the time-varying correlations separately for animate, and inanimate exemplars.

## 3. Results

### 3.1 Behavioral results

Overall, subjects performed well on the categorization task (mean accuracy 87.9%, sd 12.7) with a median RT of 457.5ms (sd 64.6). Figure 5 shows the median RT (Figure 5A) and mean accuracy (Figure 5B) separate for category (animate vs. inanimate) and stimulus condition (clear vs degraded). Analysis of variance showed that animate exemplars were easier to categorize than inanimate exemplars (compare left versus right groups in each plot), with significantly faster median RTs (*F*(1,19)=32.09, p<.0001) and higher accuracies (*F*(1,19)=6.27, p<.0001). As predicted, degrading the stimuli made the animacy categorization task more difficult (compare blue and yellow lines), significantly lowering accuracy (*F*(1,19)=41.81, p<.0001) and increasing median RT (*F*(1,19)=85.06, p<.0001) for both animate and inanimate exemplars. There was no significant interaction between animacy and degrading for either median RT (*F*(1,19)=0.04, p=0.84), or accuracy (*F*(1,19)=0.01, p=0.92).

In order to test whether distance is related to evidence accumulation, we obtained drift rates for each exemplar and subject from the LBA fits. The LBA produces a separate drift rate parameter for the correct accumulator (i.e., the animate accumulator for animate stimuli, and the inanimate accumulator for inanimate stimuli) and the incorrect accumulator (i.e., the inanimate accumulator for animate stimuli and vice versa), however only the drift rate for the correct response accumulators were included in our analysis. The resulting drift rate parameters are summarized in Figure 5C, split by animacy and degradation. Degrading the stimuli resulted in significantly slower drift rates (*F*(1,19)=41.20, p<.0001). There was a main effect of animacy on drift rate (*F*(1,19)=4.44, p<0.05), and a significant interaction between animacy and stimulus clarity on drift rate (*F*(1,19)=18.52, p<0.0001), as degrading reduced the drift rate for animate exemplars more than inanimate exemplars (see Figure 5C). In sum, we confirmed that the degraded exemplars were generally harder to categorize (in terms of longer RT, lower accuracy, and slower drift rate), demonstrating our stimulus categorization difficulty manipulation (blur) was successful.

**Figure 5.**
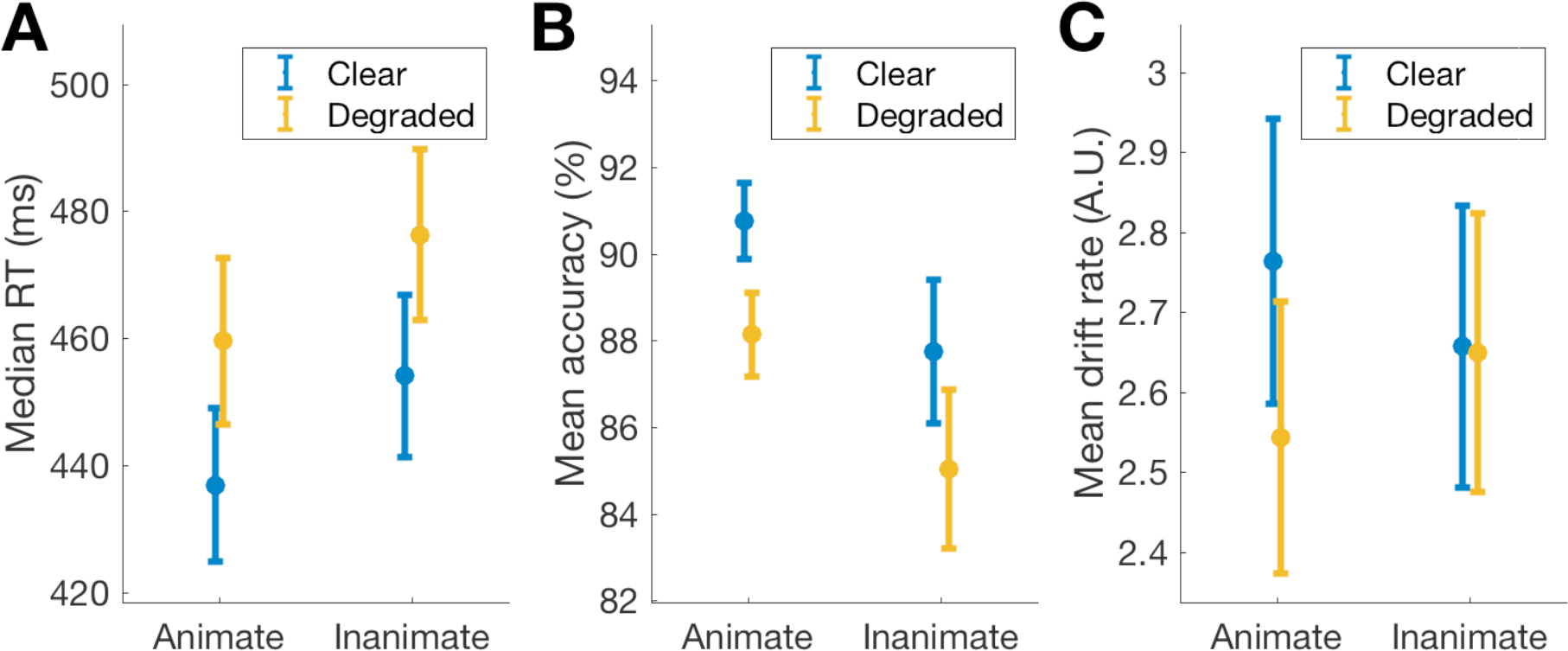
Behavioral results. The distributions of (A) the median reaction time, (B) accuracy, and (C) drift rate for the correct response accumulator, split up by exemplar category (animate/inanimate) and condition (clear/degraded, blue and orange lines). Error bars represent +/- SEM.

### 3.2 MEG Decoding

A prediction of degrading the stimuli is that decoding performance will be lower relative to that for clear stimuli. To evaluate the effect of blurring the stimuli on decoding performance over time, sliding time window classifiers were trained on predicting stimulus animacy at each time point. The results are presented in Figure 6A, as mean cross-validated decoding accuracy over subjects, separately for clear and degraded stimuli. The decoding onset in the clear condition is at 75ms, which is consistent with previous findings showing an animacy decoding onset of approximately 60 - 80 ms (Carlson et al., 2013; Ritchie et al., 2015). In the degraded condition, decoding onset was 170 ms, and decoding performance was significantly lower than in the clear condition over the entire time course (black marks above the x-axis). Peak decoding performance for degraded objects was also later, 380 ms compared to 345 ms for clear stimuli. Initially, decoding performance for the combined data for clear and degraded conditions closely matched that for the degraded condition, but then raised to a level closer to the performance in the clear condition. In sum, the neural patterns for blurred stimuli were more difficult to decode from the whole-brain MEG activation patterns than those for the clear versions of the same stimuli.

**Figure 6.**
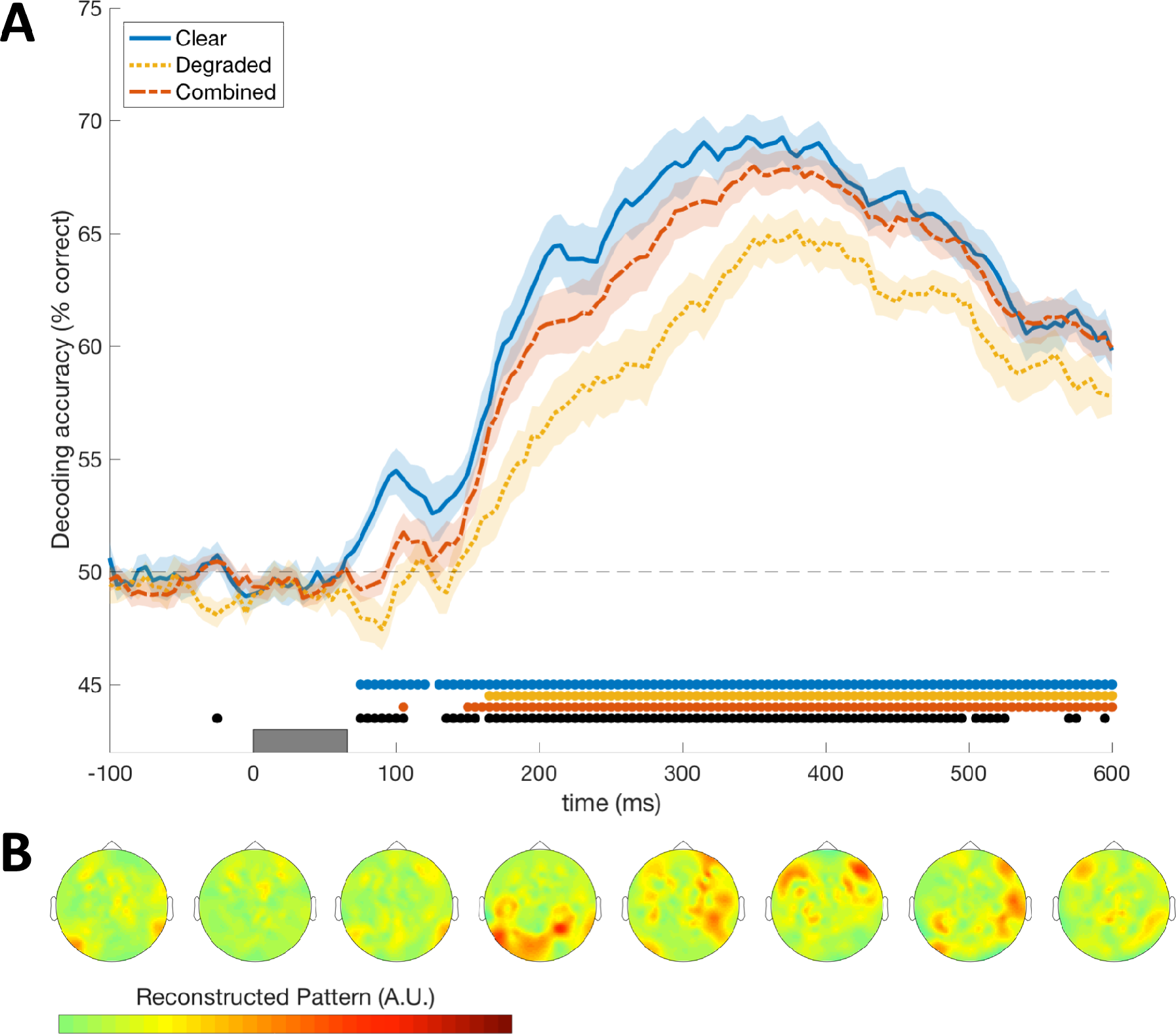
Decoding animacy from the MEG signal. (A) Decoding was performed using leave-one-exemplar-out cross-validation with 25ms sliding time window classifiers. At each time point t, the graph shows the mean classifier accuracy over subjects at the window [*t*-25ms, *t*]. Shaded areas show standard error between subjects. Colored marks above the x-axis indicate significant above-chance (50%) decoding. Black marks indicate significant (FDR-adjusted p<0.05) differences in classifier accuracy between the clear and degraded conditions. The gray bar on the x-axis indicates the time that the stimulus was on the screen (0-66ms). (B) The mean classifier weights at 100ms intervals over cross validation folds and over subjects for the combined condition. Weights were transformed into activation patterns prior to projection onto the sensors (Haufe et al., 2014).

### 3.3 Predicting behavior from representational distance

To investigate the relationship between decodability and behavior, we computed the time-varying distance to the classifier decision boundary for all the exemplars for each subject. These subject-specific distances were then rank-order correlated to the subject’s behavioral measures for each exemplar: RT, accuracy, and the drift rate as fitted by LBA. Note that for accuracy and drift rate, the RT-distance hypothesis predicts a *positive* correlation (closer to the boundary corresponds to lower accuracy and slower drift rate), however a *negative* correlation is predicted between RT and distance (closer to the boundary corresponds to longer reaction times). For ease of comparison the sign of the correlation for RT was inverted in Fig 7.

Figure 7A shows the mean time-varying correlations over subjects for drift rate (green line), RT (red line), and accuracy (purple line). The peaks of the time-varying correlations are shown in the inset bar graphs. Although drift rate appears to have an earlier and higher peak correlation than RT and accuracy, this difference was not statistically significant. The similarity between these time-varying correlations is likely due to correlations among the different behavioral measures (e.g., exemplars with fast RTs are likely to have high accuracies). These results further show that accuracy and drift rate can be predicted equally well using distance to boundary. While the LBA model has been used before to fit non-human primate neural activations (Cassey, Heathcote, & Brown, 2014), this is the first time it has been related to neuroimaging data measured with MEG. Thus, our results are promising considering that drift rate more closely represents the accumulation of evidence for a decision, compared to RT or accuracy. In sum, distance to boundary correlates with the behavioral measures as well as the fitted drift rate parameters and these follow similar trajectories.

**Figure 7.**
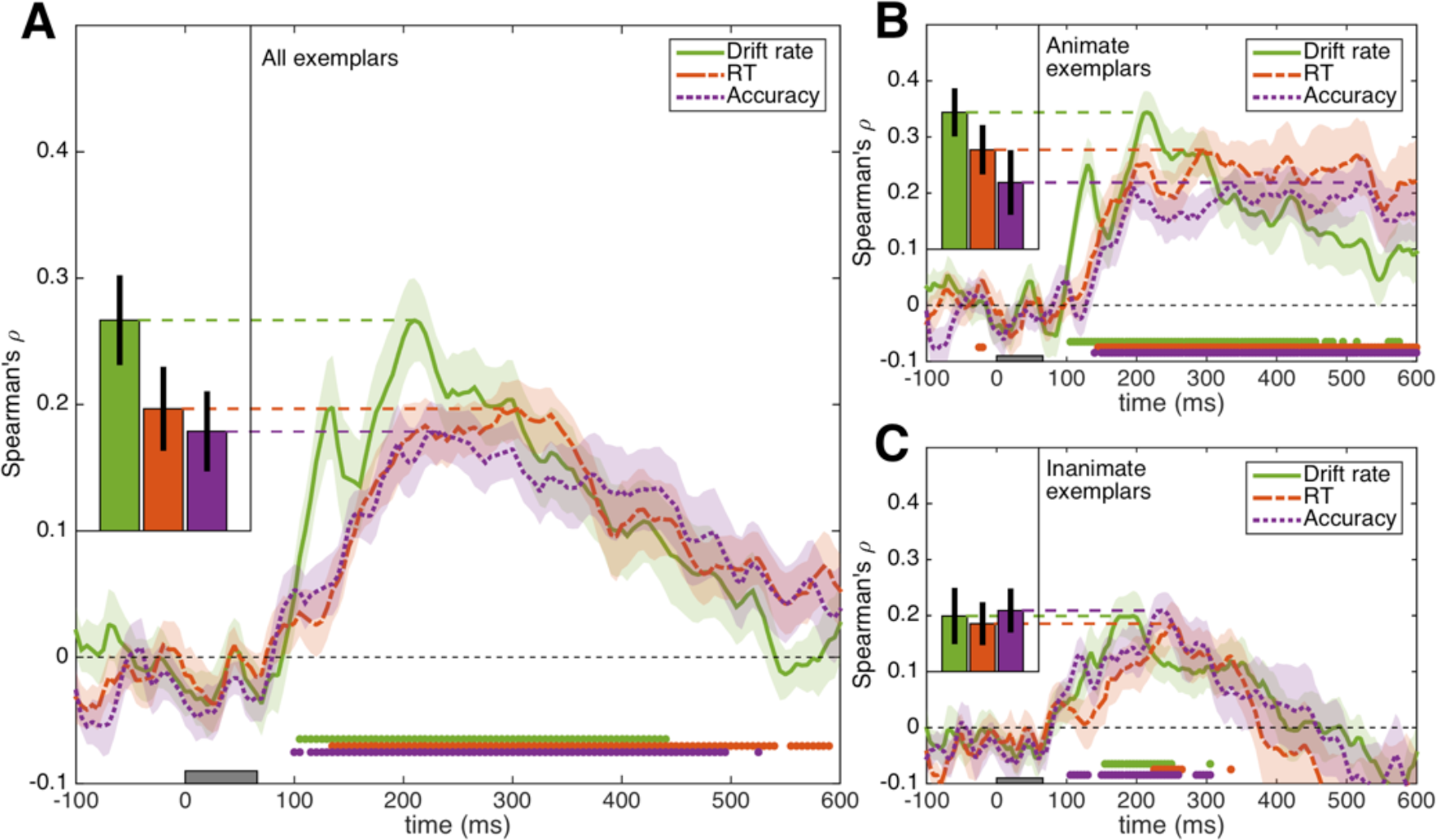
The correlation between distance to boundary, behavioral variables, and drift rate. **A.** The distance model applied to all the exemplars. **B.** Results for animate exemplars only. C. Results for inanimate exemplars. Three measures (RT, accuracy, and the drift rate parameter) were correlated with distance to boundary over time. RT refers to inverted normalized RT. Accuracy is the mean accuracy for exemplars, and drift rate was estimated by fitting LBA. Shaded areas refer to the standard error over subjects, and colored marks above the x-axis indicate significant above-zero correlations. The peak correlations with their standard errors are compared in the inset (bars top left). The gray bar on the x-axis indicates the time the stimulus was on the screen (0-66ms).

Having established that distance to boundary predicts RT, accuracy and drift rate for animacy categorization, we next investigated the relative contributions of animate and inanimate exemplars. In Ritchie et al., (2015), time-varying correlations for the inanimate exemplars were not significant. Thus the effect was specific to animate exemplars, and the same asymmetry was also reported in Carlson et al., (2014). In this study however, due to the degrading of stimuli, we also predicted a correlation for inanimate exemplars (when using both clear and degraded exemplars), as we predicted that the representations of degraded inanimate exemplars would still shift towards the classifier decision boundary (Figure 1). Figure 7 shows the result of computing the time-varying correlation separately for animates (Figure 7B) and inanimates (Figure 7C). Animate exemplars reach higher correlations on all behavioral measures, and follow the same trends as seen in Figure 7A for the combined stimulus set. In contrast, inanimate exemplars have lower correlations overall and were less sustained over time. However, significant above-chance correlations between distance and RT, accuracy, and drift rate were still present for the inanimate exemplars. In addition, more sustained correlations (more significant time points) are present for drift rate and accuracy than for RT, for the inanimate exemplars (which is of interest, as previous studies only used RT).

### 3.4 Comparing the decoding time courses with the time course of predicting behavior

Ritchie et al. (2015) found that the time-varying correlation with behavior matched the time-varying classifier decoding performance. To examine whether this relationship was present in the current study, the time-varying correlations with behavior (RT, accuracy and drift rate) were rank-order correlated to the time-varying decoding result. The results are presented in Figure 8, and show significant correlations between the results from Figure 7 and the decoding trajectory (Figure 6A), rising and falling at approximately the same time. When comparing the animate and inanimate trajectories separately (Figure 7B-C), only the animate time-varying correlation matches decoding performance. These results are consistent with the findings from Ritchie et al. (2015) who also found higher correlations between RT-correlation and decoding trajectories for the animate exemplars. In addition, the decoding trajectory also correlates significantly with the accuracy results from Figure 7, and is significant for both animate and inanimate exemplars. Correlations with the drift rate parameter are also significant, but appear lower in magnitude. This is evident when comparing Figures 6 and 7, where drift rate has a dual peak structure, which differs from the trajectories for the RT and accuracy correlations.

**Figure 8.**
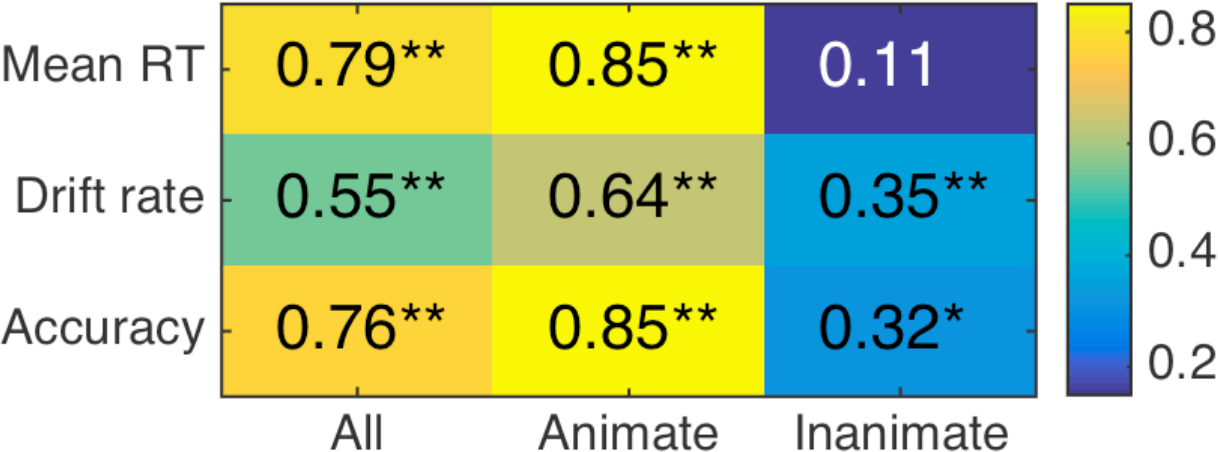
Similarity (Spearman’s p) between the decoding trajectories (Figure 6A), and correlation trajectories from Figure 7. The time varying decoding performance (for all clear and degraded stimuli combined), and time-varying correlations were rank-order correlated. High values indicate similar trajectories (e.g., matching rises, falls, and peaks). Asterisks indicate significant correlations (*=p<.01; **=p<.001).

### 3.5 Compression of degraded objects in representational space

The RT-distance hypothesis predicts that the neural patterns for degraded objects (which have slower RTs and lower accuracies; see Figure 5) will be closer to the category boundary in representational space, as the distance to the boundary represents the degree of evidence for category membership (Figure 1). The results above show that classifier accuracy is lower for degraded objects, indicating that degraded items are harder to separate in representational space (Figure 6A). Next, we showed that distance to boundary correlates with all three behavioral measures (Figure 7A), and this correlation is mostly driven by animate exemplars where it is sustained over time (Figure 7B), with some significant correlations for inanimates across a more limited portion of the time course (Figure 7C). Together, these results seem to favor our prediction based on the RT-distance hypothesis: degraded objects (both animate and, to some extent, inanimate) are closer to the boundary than their clear counterparts, and this shift in representational space correlates with RT, accuracy, and drift rate.

To visually compare our results to the predictions as shown in Figure 1A, we plotted all exemplars in both clear and degraded states relative to the decision boundary in representational space (at the time of peak correlation between drift rate and distance to boundary (210ms), Figure 7A) in Figure 9. On the Y-axis, the stimuli are ordered by their typicality ratings. Comparing Figure 1A and Figure 9 suggests that the prediction that degraded stimuli are located closer to the category boundary than clear stimuli holds only for animate, but not inanimate, exemplars. This result is consistent with previous results from Ritchie et al. (2015) and Carlson et al., (2014) which showed no clear relationship between distance to boundary and reaction time for inanimate stimuli. Although, even if the basic relationship between behavior and distance to the classifier boundary does not hold for inanimate exemplars, it is still surprising that there is also no relationship between degrading inanimate stimuli and where they are located relative to the category boundary. This asymmetric compression of representational space is however consistent with the interaction between animacy and degrading on the fitted drift rates (Figure 5C), which suggests that there is little or no effect of degrading on the representation of inanimate exemplars.

**Figure 9.**
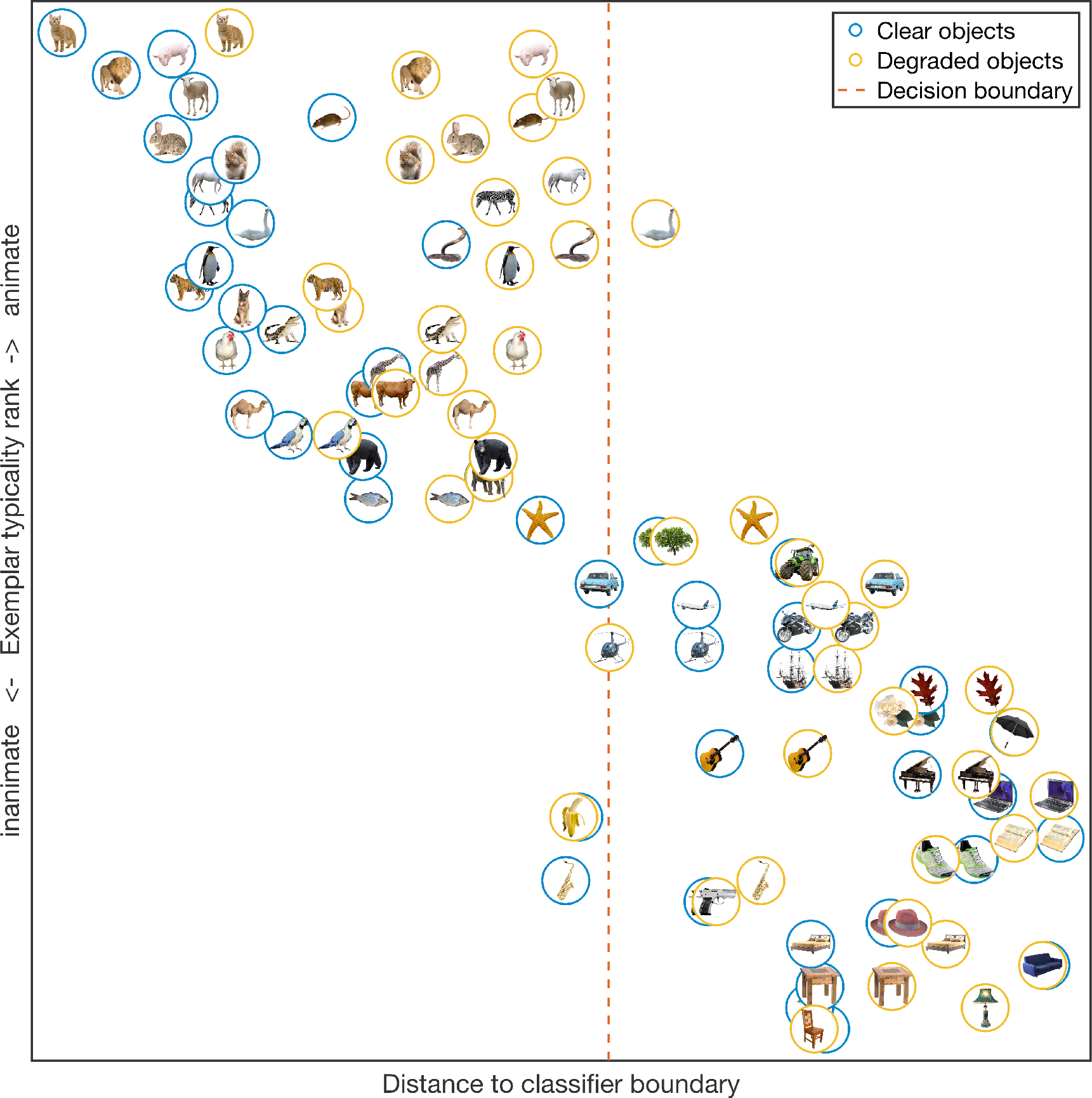
Reconstruction of the representational space for animacy decoding. The location of exemplars in their clear state are plotted in blue circles, and their degraded counterparts in yellow circles. The x-axis represents distance from an object to the boundary, and is the mean distance to the boundary over all subjects at 210ms. The exemplars are ordered on the y-axis according to their typicality rating obtained from 40 participants on MTurk. Note that some objects (e.g., the degraded starfish) have a negative mean distance, and are thus placed on the opposite side of the boundary. Note that degraded animate exemplar representations are in general closer to the linear decision boundary than the clear versions, demonstrating compression of representational space for animate but not inanimate objects.

The compression of exemplar distances towards the animacy boundary is also predicted to match slower drift rates (as shown in Figure 1B). To compare our results to this second prediction, we plotted the mean drift rates over subjects against the mean distance to boundary over subjects for each exemplar in both states in Figure 10 separately for animate (Figure 10A) and inanimate (Figure 10B) exemplars. The result is shown at the time of peak correlation between drift rate and distance to boundary (210ms, Figure 7A). The exemplars are plotted as a function of their mean distance to boundary (x-axis), and mean drift rate (y-axis). The RT-distance hypothesis predicts that exemplars with a higher drift rate are located further away from the boundary. Lines in Figure 10 connect the two states of each exemplar and are colored green if this prediction is true for each exemplar. This is the case for most of the animate exemplars (Figure 10A), but (not surprisingly, considering the asymmetry in Figure 9) for only a few of the inanimate exemplars (Figure 10B).

**Figure 10.**
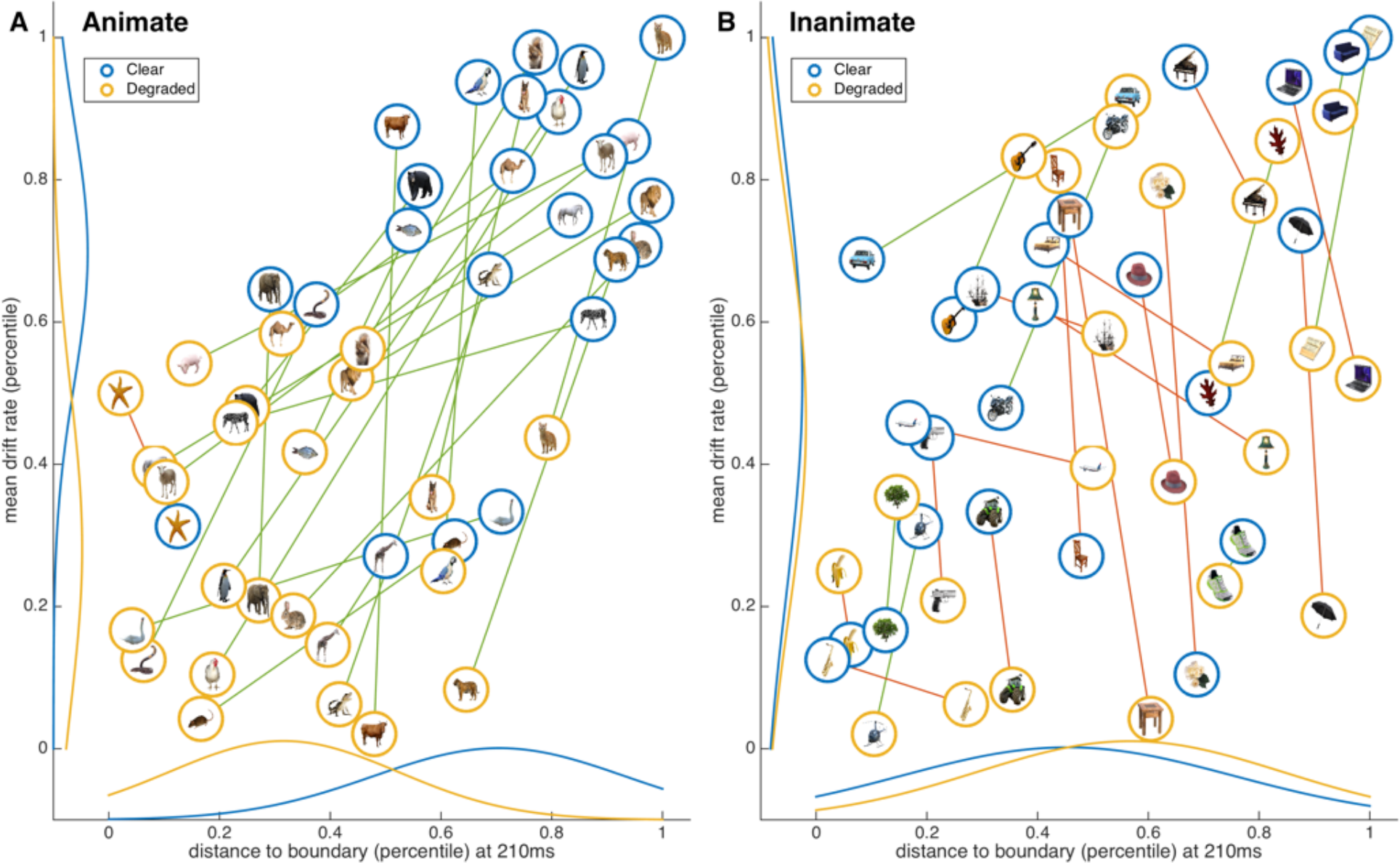
The effect of degrading on the relationship between drift rate and distance to boundary for (A) animate and (B) inanimate exemplars. Items were rank-ordered for mean drift rate over subjects, and mean distance to boundary at the peak LBA prediction time (210ms, see Figure 7A). Here, the y-axis represents the decision boundary. Blue circles are the exemplars in their clear versions, and yellow circles degraded versions. Lines connect the two versions of each exemplar, and are colored green if the RT-distance hypothesis correctly predicts the direction of the relationship (i.e., the degraded versions of the objects are closer to the boundary and have a slower drift rate), and red if not. The distributions of the exemplars on the variables are shown on the axes.

## 4. Discussion

Our aim was to test the RT-distance hypothesis in the context of a specific prediction, namely that degrading objects will result in a compression of representational space, and that the shift in representational space for degraded objects will match impaired categorization behavior. The results showed that degraded object images produced slower RTs, lower accuracies, and slower evidence accumulation rates in an animacy categorization task (Figure 5). When examining the distances of individual objects, we observed an asymmetric compression: only the degraded animate objects had moved closer towards the classifier boundary. Unexpectedly, there was not a corresponding consistent shift towards the boundary for degraded inanimate exemplars. In addition, we found that distance to boundary correlates with drift rate, which is a measure that is more closely related to the decision process than descriptive statistics (mean RT or accuracy).

In this study, all MEG channels were included for training and testing the classifier, as is standard for many MEG decoding studies (e.g., Carlson et al., 2013; Cichy et al., 2014; Grootswagers et al., 2017; Kaiser, Azzalini, & Peelen, 2016). A limitation of this approach is that the spatial source of the decodable signal is unknown. Because participants were performing an animacy task, it is not possible to entirely exclude a contribution from the decision process (e.g., originating in frontal executive areas) as a possible source of the decodable signal. However, previous research has found no difference in decoding performance between active animacy categorization versus a distractor task (Ritchie et al., 2015). In addition, the representational structure in the MEG response to visual objects has been found to correspond best to fMRI representations in the ventral visual stream, showing that the early and late MEG response following stimulus onset matches the fMRI responses in V1 and IT respectively (Cichy et al., 2014, 2016). Moreover, examining the classifier weights (Figure 6B) suggests the decodable signal originates from occipital sensors at 200ms, and more temporal at 300ms. This suggests that the most likely sources of the decodable signal in our study originate from areas in the ventral visual stream.

### 4.1 Asymmetric effects of stimulus degradation on the neural representation of animacy

We found asymmetrical effects for animate and inanimate objects. Correlations between distance and behavior (RT/accuracy/drift rate) in this study were driven by animate stimuli. Inanimate stimuli had smaller and less sustained correlations between distance and behavior, and did not show compression towards the classifier boundary in representational space for degraded versions of the stimuli. Previous studies also found that correlations between distance and RT were almost exclusively driven by animate stimuli (Carlson et al., 2014; Ritchie et al., 2015), which is consistent with our results. Our study differed in many aspects from earlier studies, for example, by using gray scale stimuli on a controlled background, and excluding human or human face stimuli. Human faces/bodies have faster RTs (Crouzet, Kirchner, & Thorpe, 2010) and highly decodable neural responses (Carlson et al., 2013; Kaneshiro et al., 2015; Kriegeskorte et al., 2008) and could therefore potentially significantly drive the RT-distance correlations in previous studies. However, we have shown these findings are robust when human faces/bodies are omitted (also note that correlations were calculated within-subject, rather than on the pooled group means as in previous work). Carlson et al. (2014) argue for a conceptual difference between the two animacy categories, suggesting that ‘inanimate’ is not an equivalent category to animate, for example, because inanimate is negatively defined (i.e., as ‘not animate’) and is less restricted than the animate category. Furthermore, a clear hierarchical subdivision (e.g., vertebrate - invertebrate) has been reported only within the animate category (Kiani et al., 2007).

We observed that degraded versions of animate objects had compressed towards the boundary, consistent with the RT-distance hypothesis. The lack of compression for the inanimate side of the boundary, and the absence of strong correlations between inanimate categorization behavior and distance to boundary, suggests that even though the animate/inanimate distinction is highly decodable, it does not sufficiently capture the structure of brain representations linked to responding ‘inanimate’. Future work could explore this issue using a different categorization task where both categories are similarly constrained (e.g., faces versus tools), or use a different model of the animacy task (e.g., model it as an animal detection task, instead of animacy categorization). However, it is still possible that responses to other category dichotomies will be based on one category versus ‘not’ that category. For example a processing bias for faces means they are easier to recognize than tools (cf. Wu, Crouzet, Thorpe, & Fabre-Thorpe, 2015) and an effective face/tool categorization strategy would be to simply try to detect whether or not a stimulus is a face. In addition, outside the lab, objects are categorized effectively without having to exhaustively test all two-way categorization combinations. Thus, instead of using dichotomous categorization, applying a different task to examine the RT-distance relationship might yield new insights, for example by using go/no-go tasks (Crouzet et al., 2010; Kirchner & Thorpe, 2006; Thorpe, Fize, & Marlot, 1996).

The amount of compression towards the decision boundary was different between individual exemplars (Figure 9). Even though the degraded stimuli were equated for recognizability in an object-naming task, some showed a larger displacement towards the boundary than others (e.g., compare difference between clear and degraded versions of the fish and the sheep in Figure 9). The naming task may be more difficult than the categorization task, and although they likely rely on the same underlying representation (Riesenhuber & Poggio, 2000), different amounts of evidence may be required for naming versus categorization. For example, some animals might be equally easy to name, but when making an animacy decision, some animals are more typically animate than others, which likely makes them easier to categorize than less typical animals. Typicality ratings for our stimuli indeed increased with distance to boundary, but again only for the animate stimuli (Figure 9). Animacy categorization is known to be influenced by typicality (Posner & Keele, 1968; Rosch, 1973; Rosch & Mervis, 1975). Typical exemplars have also been found to be more decodable (Iordan, Greene, Beck, & Fei-Fei, 2016). For example, mammals such as the cat, tiger, and squirrel are all far from the boundary and have matching fast reaction times and high drift rates. Conversely, the fish is closer to the boundary, and is possibly less typically conceived of as ‘animate’, for example, because fish move and behave differently than mammals. An extreme example is the starfish, which is the animal closest to the boundary in the clear condition, and is on the wrong side of the boundary (i.e., consistently predicted ‘inanimate’ by the classifier) in the degraded condition, suggesting that it is hard to categorize it as animate (see Figure 9). We found that subjects mostly categorized the starfish as inanimate, and that, when asked, some reported that they do not consider it an animate object. Note that in order to observe a correlation between distance to the boundary and reaction times, distances and reaction times have to systematically differ between exemplars in the same category. Our results thus provide some support for the presence of a more continuous than dichotomous neural representation of animacy in the brain (cf. Connolly et al., 2012; Sha et al., 2015).

Further evidence in support of an animacy asymmetry comes from our observation that inanimate stimuli in general needed to be blurred more to equate their recognizability to animate exemplars (see section 2). This suggests that animate objects are more homogenous than inanimate objects, thus there is greater variance in the ‘inanimate’ than ‘animate’ category (which could be caused by the lack of categorical structure in inanimates). This is consistent with animate objects in general sharing more features than inanimate objects (Garrard, Ralph, Hodges, & Patterson, 2001; McRae, de Sa, & Seidenberg, 1997). The blur filter removes details from the stimuli, but object shape is preserved. It could be the case that the general shape within the broad category of animate objects is more similar, and provides more alternatives (cf. Bracci & op de Beeck, 2016). Even though some higher-level animal subdivisions might be visually dissimilar (e.g., compare fish with birds), subgroups often share similar shapes. For example, outside the laboratory, in order to recognize a blurry zebra (e.g., without glasses), likely alternatives that have similar shapes (e.g., horse, deer, moose) need to be eliminated. In contrast, viewing a blurry piano provides less alternatives which share the same shape. Together, this supports the notion that inanimate and animate are not equivalent categories, which is consistent with patient studies that found selective deficits in recognition of animate or inanimate objects (Caramazza & Mahon, 2003; Caramazza & Shelton, 1998). Future research could explore whether homogeneity of the category determines the effect of degrading stimuli, by using the distance to bound approach with more homogenous groups of inanimate objects, such as tools or fruits, or restricting the animate category to only domestic animals.

### 4.2 Drift rate is predicted by distance to boundary

The drift rate parameter from the LBA model more directly reflects the speed of evidence accumulation, and builds upon previous work that used RT as a proxy for the decision process (Carlson et al., 2014; Ritchie et al., 2015). We found that correlations with LBA drift rate were on par with those for RT, and that there were some qualitative differences in the trajectories. The correlation between LBA drift rate and distance peaked earlier than RT, and had an earlier onset—although this should be interpreted with caution as earlier onsets can be caused by stronger signal-to-noise rather than true underlying differences (Grootswagers et al., 2017). Moreover, drift rate followed a different trajectory than the category decoding trace (as seen in Figure 8), which suggests it may capture a different part of the (neural) decision process. It is interesting that the drift rate trajectory does have a dual peak structure, which resembles previous results of MEG animacy decoding (Carlson et al., 2013; Cichy et al., 2014; Ritchie et al., 2015), but which was not present in our MEG decoding results. This dual-peak structure may suggest that at some point (in the time period between the peaks), distance to boundary is used to a lesser extent for forming the categorization decision. Alternatively, the decision could have already been made after the first peak, as categorization happens very fast (Crouzet et al., 2010; Kirchner & Thorpe, 2006; Thorpe et al., 1996), and the second peak reflects, for example, a feedback process that has the same representational structure (i.e., the same distances). Taken together, it is sensible to relate distance directly to evidence accumulation using drift rate as it is more closely related to decision processes and therefore it may successfully relate to a wider range of behavior than RT alone. Future research could explore this further by using tasks that allow more for more variance in RT and accuracy (e.g., incentivizing different speed-accuracy trade-offs). Moreover, as we showed it is possible to link one decision model with neural distance to boundary, this could be tested with other models of decision making, such as exemplar-based models of choice (Nosofsky & Stanton, 2005) or their LBA-based extension (Donkin & Nosofsky, 2012).

### 4.3 The neural dynamics of visual object categorization

We found that significant decoding performance of clear stimuli started around 70ms, and peaked at 345 msec, occurring later for degraded stimuli (165 ms and 380 ms, respectively). Note that differences in decoding onsets have to be interpreted with caution, as a later onset can be a result of lower overall decoding performance, or differences in variance (Grootswagers et al., 2017; Isik et al., 2014). Still, we observed that the first local maximum in the decoding trace for the clear objects was absent for the degraded objects (Figure 6A). This difference suggests that some information in the early response is predictive of animacy in the clear condition. As the timing of the early peak corresponds to early to mid-level visual areas (Cichy et al., 2016; Thorpe et al., 1996; VanRullen & Thorpe, 2001), the predictive information in the first local maximum in the clear decoding condition could reflect low-level visual information (e.g., high spatial frequencies) that is removed by degrading the stimuli (Kirchner & Thorpe, 2006).

We found that the peak correlation between distance and drift rate occurred at 210 ms, and the onset of significant correlations with drift rate was at 100 ms (Figure 7). These values did not match the onset or peak of decoding, suggesting that the optimal time for read out does not necessarily correspond with the time that the information can best decoded from the signal. In contrast, Ritchie et al., (2015) found correlations during the whole time period of significant decoding. A possible explanation for this difference is that the fast-paced task and short stimulus duration (66 ms compared to 500 ms in Ritchie et al., (2015)) in the current study may have promoted a faster read out of animacy by exploiting low level visual cues (Hong, Yamins, Majaj, & DiCarlo, 2016; Kirchner & Thorpe, 2006; Thorpe et al., 1996). This would in addition explain why exemplars such as the banana and helicopter, which have more rounded shapes than other inanimate objects, are closer to the boundary (Figure 9).

### 4.4 Conclusion

In this study we tested whether representational space is compressed when degrading stimuli, and whether this matches behavior in a dichotomous categorization task. We found that degrading stimuli made them harder to categorize, and that this was accompanied by a compression of representational space, as predicted by the RT-distance hypothesis. This compression was only observed for animate stimuli, suggesting an asymmetry in in the neural representation of animacy. Moreover, we showed that neural distance to boundary can be directly related to a current model of evidence accumulation (LBA) as the fitted drift rates from this model correlated with distance to the boundary. Connecting linear classifiers to models of the decision processes is a step towards relating brain imaging to behavior, a fundamental and complex challenge in cognitive neuroscience (de-Wit et al., 2016; Forstmann & Wagenmakers, 2015; Purcell & Palmeri, 2016).

## Acknowledgements

This research was supported by an Australian Research Council (ARC) Future Fellowship (FT120100816) and ARC Discovery project (DP160101300) awarded to T.A.C. S.G.W. is supported by an Australian NHMRC Early Career Fellowship (APP1072245). The authors declare no competing financial interests.

